# Insights into a viral motor: the structure of the HK97 packaging termination assembly

**DOI:** 10.1101/2023.02.24.529869

**Authors:** Dorothy E.D.P. Hawkins, Oliver Bayfield, Herman K.H. Fung, Daniel N Grba, Alexis Huet, James F. Conway, Alfred A. Antson

## Abstract

Double-stranded DNA viruses utilise machinery, made of terminase proteins, to package viral DNA into the capsid. For *cos* bacteriophage, a defined signal, recognised by small terminase, flanks each genome unit. Here we present the first structural data for a *cos* virus DNA packaging motor, assembled from the bacteriophage HK97 terminase proteins, procapsids encompassing the portal protein, and DNA containing a *cos* site. The cryo-EM structure is consistent with the packaging termination state adopted after DNA cleavage, with DNA density within the large terminase assembly ending abruptly at the portal protein entrance. Retention of the large terminase complex after cleavage of the short DNA substrate suggests that motor dissociation from the capsid requires headful pressure, in common with *pac* viruses. Interestingly, the clip domain of the 12-subunit portal protein does not adhere to C_12_ symmetry, indicating asymmetry induced by binding of the large terminase/DNA. The motor assembly is also highly asymmetric, showing a ring of 5 large terminase monomers, tilted against the portal. Variable degrees of extension between N- and C-terminal domains of individual subunits suggest a mechanism of DNA translocation driven by inter-domain contraction and relaxation.

## INTRODUCTION

Encapsulation of the genome represents a key step in viral assembly and life cycle. For double stranded (ds) DNA bacteriophage, the genome is packaged into a preformed protein shell, or capsid, through the dodecameric portal protein, which acts as a door (1). Packaging often produces expansion of the capsid from an immature form, known as a prohead, to a larger, fully mature state.

Translocation of DNA against mounting internal pressure requires ATPase activity, provided by of a ring of five large terminase subunits (2–6). Each monomer constitutes an N-terminal ATPase (NTD) (28) and C-terminal endonuclease domain (CTD) adjoined by a linker. This large terminase motor is thought to reach forces of up to 100 pN (7–9) and thus represents the most powerful biological machine studied. Large terminase functions akin to other ring-shaped, oligomeric translocases, which also utilise ATP hydrolysis cycles to power translocation through the central pore (10). Most dsDNA bacteriophage also employ small terminase proteins for recognition of the viral genome (11–14). Small terminases have also been shown to stimulate large terminase packaging activity (15–17).

Although mechanisms of packaging are broadly conserved among the dsDNA phages, further differentiations can be highlighted based on processing of the viral DNA at the initiation and termination of packaging. *Cos* viruses, such as λ, P2, and HK97—the subject of this paper— package exact genome-length DNA units (18,19), separated by consecutive *cos* (cohesive) sites in a genome concatemer (20). For *pac* viruses, including P22, P1, SPP1 and T4, packaging is initiated from a *pac* (packaging) site within the genome (21–23) and terminates in response to ‘headful’ pressure within the prohead, leading to encapsulated genomes varying in length from ~103–110% (24). Finally, Φ29-like phages produce unit-length DNA genome copies, where each 5’ end is covalently bound to viral protein gp3 (25) which facilitates packaging.

Portal proteins display highly divergent sequences and molecular weights, but *in situ* within the viral capsid, consistently present as dodecameric rings (26–28). The monomeric portal protein structures share conserved protein folds and domain arrangement comprising the clip, stem, wing, and crown domains. Positioned at the base of the portal protein, the clip domain interacts with the large terminase motor during assembly, and later the adaptor proteins for tail attachment (26). The stem contains the “tunnel” a-helix that lines the internal channel with several negatively charged residues (29). The most structurally divergent domain, the wing, coordinates contact with the capsid proteins (26–28). Lastly the crown, exposed at the inner surface of the capsid, interacts with packaged DNA (30).

The portal protein displays remarkable plasticity, proposed to facilitate symmetry-mismatching interactions with the capsid and terminase. This flexibility also permits propagation of pressure changes within the capsid to the terminase (31), in order to induce packaging termination. Indeed, the DNA in mature phage P22, is spooled tightly around the portal in an arrangement which appears to be incompatible with the procapsid portal form (31). Meanwhile several discrete mutations within the P22 and *λ* portal cores produce over-packing phenotypes (32,33), indicating unsuccessful termination. Conformational change of the portal protein during DNA packaging appears conserved among dsDNA viruses, with marked variability between procapsid and mature head portal structures for T7, Φ29, p22 and P23-45 (30,31,34–38).

Much mechanistic understanding of the large terminase motor has been drawn from single molecule optical tweezer studies on the phage Φ29. The Φ29 large terminase mechano-chemical cycle has been shown to alternate between two phases: During the dwell phase ATP binds cooperatively to each subunit around the ATPase ring, and the DNA substrate remains stationary. During the burst phase, DNA translocation into the capsid occurs in 4 subsequent 2.5 base pair steps corresponding to 4 ATP hydrolysis events (39). A cryo-EM reconstruction of the intact Φ29 packaging motor, comprising the capsid, pRNA, large terminase and DNA, revealed the five ATPase domains, in a “cracked” helical conformation, stabilised by ATP*γ*S (3) This conformation contrasts planar structures seen for the apo Φ29 ATPase (33,34) and the ADP-bound form of highly related phage ascc-φ28 large terminase (40). The transition between the cracked helical and planar states likely represents the burst phase, which has inspired a unique DNA translocation model (3,40). However, other terminase-packaging systems may well vary wildly from the Φ29 model, which unusually utilises pRNA and does not require small terminase proteins (41,42).

Whilst a wealth of structural data comprising individual packaging proteins has emerged for dsDNA viruses, the field is only starting to move towards analysing complete packaging systems. For bacteriophage HK97, the subject of current study, structural information is so far available only for separate components including large and small terminase proteins (43) and prohead II (the immature prohead state at packaging initiation) (44). The X-ray structure of monomeric large terminase revealed the classical ATPase and nuclease domains adjoined by a short linker (43,45). In common with most other phage, the small terminase contains 9 subunits arranged in a circular structure exposing N-terminal HTH motifs around the outside of the molecule, where they form a positively charged rim (36,38).

The cryo-EM structure of the active HK97 packaging machinery presented here, in combination with previous work establishing a functional packaging assay (43), begins to shed light on a number of unanswered questions for *cos* phage: (i) how do these viruses overcome the symmetry mismatch between the 12-subunit portal and pentameric large terminase, (ii) does small terminase remain bound throughout packaging and (iii) what is the mechanism of coordinated ATP hydrolysis around the large terminase ring and how are the chemical events of ATP hydrolysis coupled to mechanical translocation of DNA.

## MATERIAL AND METHODS

### Packaging assays

Individual packaging protein components of the packaging assays were expressed and purified as described (43). The DNA substrate used represented a linearised pUC18 plasmid with an engineered *cos* site from −312 to +472 of the cleavage sites. Limiting the packaging time from 30 minutes to 2 minutes significantly improved large terminase retention, and the addition of DNAase1 to the final sample improved the background signal.

### Data collection

R 3.5/1 Quantifoil, with 2 nm Ultrathin Carbon, 200 mesh copper grids were glow discharged for 60 seconds at 15 mA in a PELCO easiGlow™. A 3 μl sample was prepped immediately prior to vitrifying using the Vitribot IV, with blot force −5 and blot time 2 seconds (54). This grid was subject to a 72-hour data collection at the UK National eBIC (Electron Bio-Imaging Centre) at the Diamond Light Source (Harwell Science and Innovation Campus) on an FEI Titan Krios instrument using a K3 detector. The data collection parameters are summarised in **Table S1**.

### Reconstruction of the prohead/portal/motor complex (Fig S2)

Data were processed in RELION 3.1 (46). Motion correction was performed using MotionCor2 (47,48) and corrected micrographs were then subject to CTF estimation using CTFFIND 4 (49). Optimised picking parameters were achieved using Topaz, as an external RELION job (50). A total of 352,600 particles were picked and subject to 2D classification. The best classes were selected, and duplicates removed, leaving 82,279 particles for icosahedral refinement. This map was subject to post processing and CTF refinement before a second round of refinement (51) (**Fig S2 A**). The resolution of the final post processed reconstruction is 3.06 Å (FSC = 0.143).

During icosahedral refinement signal from asymmetric features of the virus is averaged out evenly over the map. Thus, in order to model the portal and motor, the icosahedral 3D reconstruction of the combined data sets was first subject to RELION symmetry expansion (51)(in symmetry point group I3), which generates a 60-fold increased set of particles. Knowing the location of the vertices, we could determine new extraction coordinates (52) for re-extraction of each vertex whilst binning to 5.2 Å/pixel. This produced 60 subparticles per capsid particle. Subparticles were subject to 3D classification without alignment to separate subparticles containing portal and motor signal, from pentameric capsomers. Initially a cylindrical mask, created using relion_helix_toolbox (53) with a soft edge of 2 pixels and extension of 2 pixels, was used to mask out contributing signal from the capsid, centred on the expected position for the portal and motor. One clear portal-containing class emerged, which was used as a template for a tighter mask applied to the original subparticle set for further classification.

To reconstruct the portal protein structure, particles from the portal-motor class were selected and subjected to 3D classification without performing image alignment. The classes displaying the most defined secondary structure were selected and re-extracted into a smaller box encompassing only the portal (1.34 Å/pixel). These 57,279 particles were refined using C12 symmetry. The 3D map was then subject to postprocessing reaching 2.98 Å resolution (FSC = 0.143) and local resolution estimation to calculate resolution within different local sections of the map (**Fig S2 B**).

The motor signal was isolated by re-extraction of particles at the portal vertex into a smaller box size encompassing the motor density only. These were subject to 3D classification without performing image alignment, and classes displaying clear individual subunits and domains were subject to a second classification into a single class. This was used as a template for a mask used for particle subtraction. This class contained just 17,584 particles. Subtracted particles were subsequently subject to refinement with local angular searches (1.8 °) followed by post-processing reaching a final resolution of 8.8 Å (FSC = 1.43) (**Fig S2 C**).

The run_data.star file from the motor refinement was used as a template for re-extraction to include portal density. The same protocol of particle subtraction was employed, using a mask to encompass the whole portal/motor complex. Subtracted particles were then refined with limited angular searches of 1.8 °. After postprocessing the resolution of the portal/large terminase complex reached 7.4 Å (**Fig S2 C**). The portal/motor complex was then reextracted to include the entire prohead and subject to symmetry expansion in C_12_. Particles were subject to 3D classification without alignment and the best 5 classes selected. Duplicates were removed leaving 8354 particles for 3D refinement (**Fig S2 C**). The resolution of the complete packaging complex was 8.3 Å (**Fig S2 C**).

### Model building

Portal and prohead atomic models were built in Coot 0.8.9.3 (54) and refined using Phenix 1.19 Cryo EM Real Space Refinement (55). For the asymmetric prohead unit model PDB model 3E8K (44) was used as a starting model (44) and each of the 7 chains refined separately from residues 130 – 382. The first 103 residues are cleaved off in the transition from Prohead I, and density for residues 104 to 129 was not sufficiently defined for model building suggesting flexibility (56). For the portal protein, a starting model for a single chain was established using an AlphaFold prediction (57) using uniprot reference P49859. This was symmetrised 12-fold, and residues 32 - 398 refined into the C_12_ map. Model validation parameters are summarised in Supplementary **Table S3**.

## RESULTS

### Stalling and stabilising the DNA packaging motor

DNA protection assays were utilised to examine optimal stalling conditions for the HK97 large terminase DNA packaging motor. This was deemed desirable in order to ‘fix’ the DNA packaging motor in a single conformation, thus preventing unsynchronised packaging and motor detachment. The presence of a *cos* site within the substrate DNA, as well as small terminase, were critical, and the ideal ATP concentration for stalling was pinpointed to 75 μM (**Fig. 1 A**). The optimised sample was then flooded with 5 mM ATP*γ*S and visualised by negative stain electron microscopy (**Fig. 1 B**) to check for associated packaging motors. Cryo-electron microscopy was then used to obtain higher resolution images (**Fig. 1 C**), followed by single-particle analysis. An asymmetric reconstruction comprising prohead II and the portal/large terminase complex, was derived at 8.3 Å. Despite limited resolution, the reconstruction contained well defined density at the unique portal vertex, likely corresponding to a large terminase oligomer. Local refinement of each protein constituent is described subsequently.

**Figure 1:**
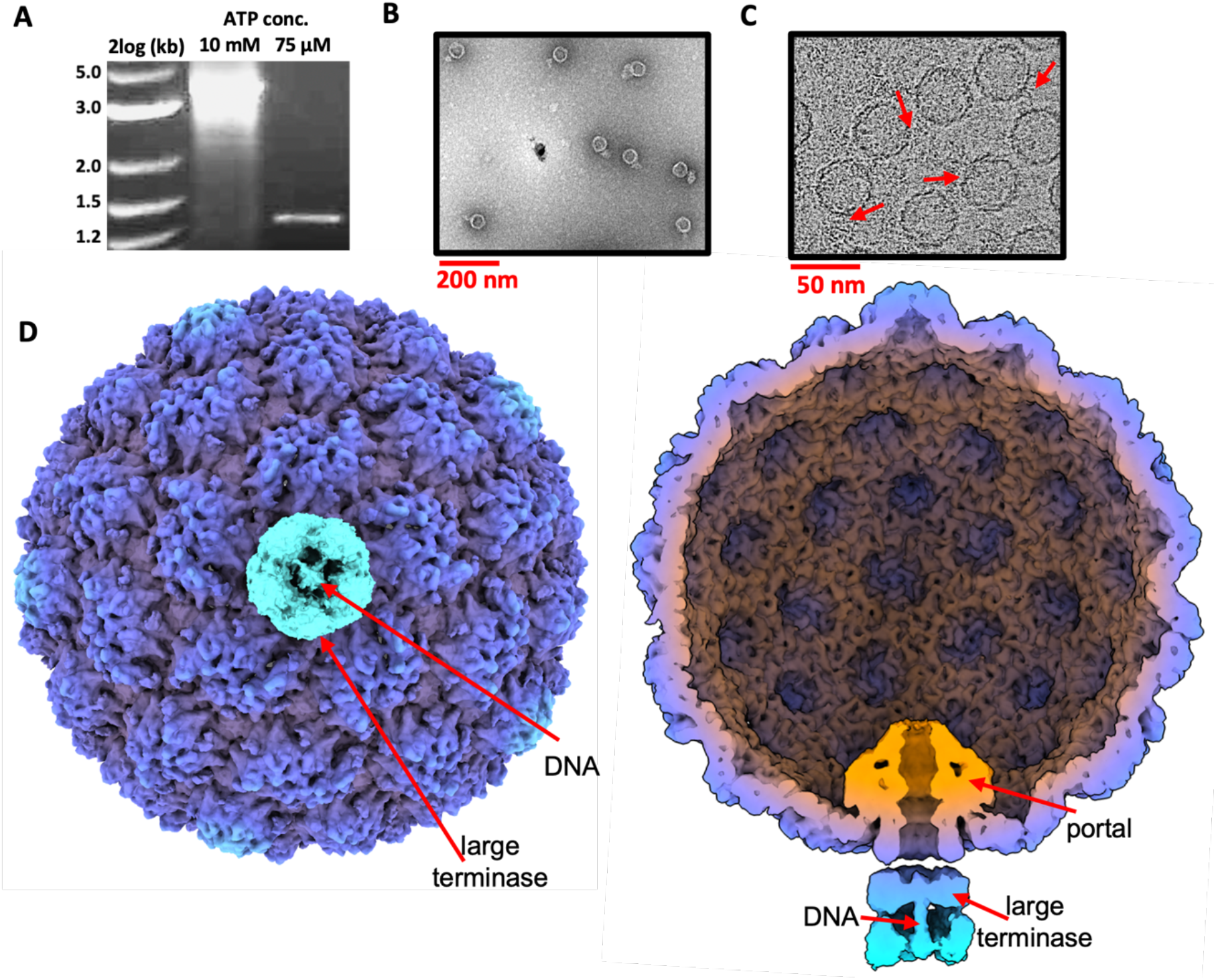
Stalled HK97 packaging complexes. **A** sample visualized by negative stain electron microscopy. **B** cryo-EM screening of sample prepared using R 3.5/1 grids with 2 nm Ultrathin Carbon film. Red arrows highlight obvious DNA packaging centres. **C** asymmetric reconstruction of the complex viewed from the motor. **D** external and cross-sectional views of the asymmetric reconstruction.

### Structure of HK97 Prohead II

The cryo-EM structure of the prohead was determined at 3.06 Å using icosahedral averaging (**Fig. 2**). EM density (**Fig. 2 C**) allowed for refinement of the previously determined 3.6 Å crystal structure (44). The prohead measures 550 Å in diameter and shows dislocated or ‘skewed’ trimers within the hexamers (**Fig. 2 A**). This is consistent with structures of prohead II (44), depicting a stage of capsid maturation prior to expansion and after cleavage of the scaffolding domains from the major capsid protein. This indicates that proheads remain unexpanded during packaging and stalling. Importantly, this model, determined for proheads containing the packaging machinery, represents a more biologically relevant form than reported in the crystal structure (44).

**Figure 2:**
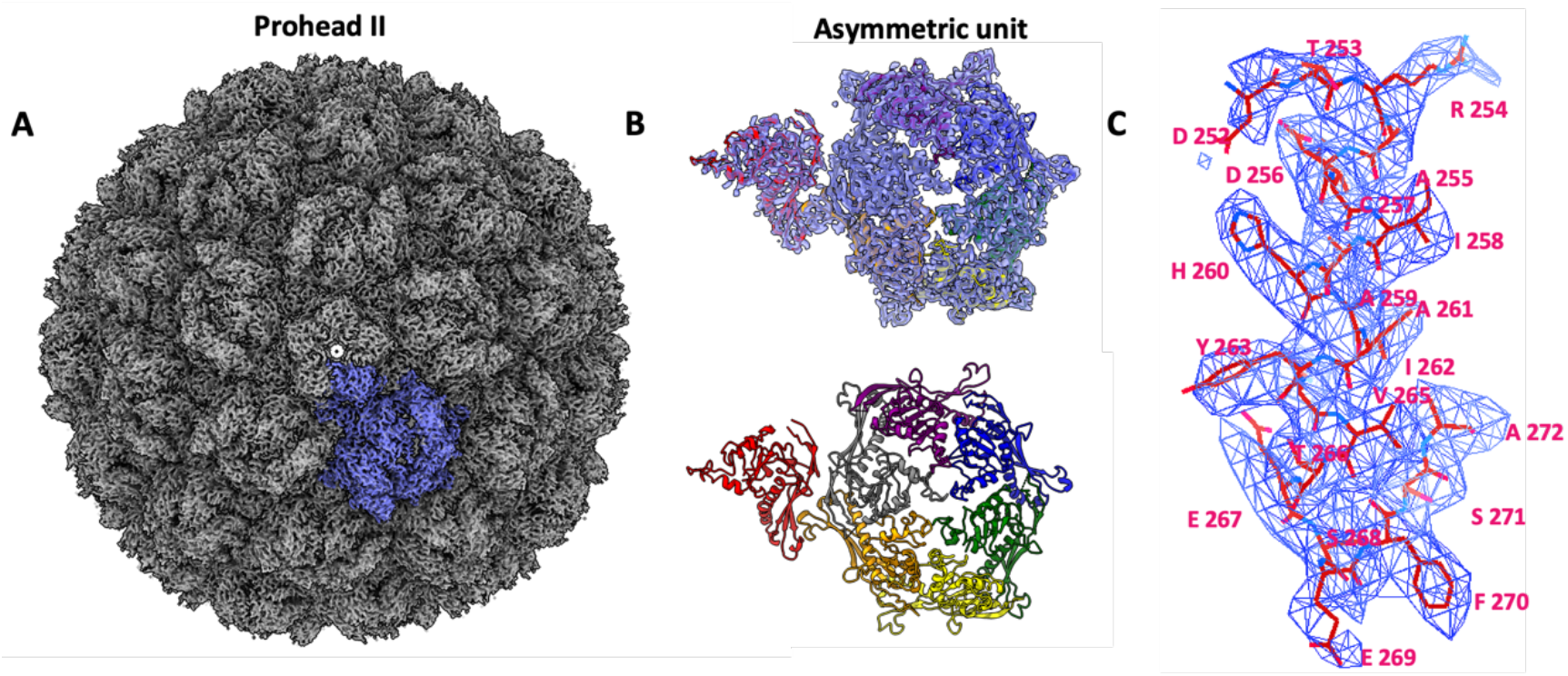
Icosahedral reconstruction of the HK97 prohead II at 3.06 Å resolution. **A** - EM density viewed along the five-fold axis. **B** - ribbon diagram of the asymmetric capsomere unit shown alone (bottom) and fit into the corresponding density map (top). **C** - representative EM map shown for a prohead segment fitted with corresponding atomic model.

### Structure of HK97 portal protein

The portal protein reconstruction shows clear 12-fold symmetry with an overall “mushroom” shaped architecture (**Fig. 3**). An atomic model was built into the 2.98 Å resolution map. The clip, stem, and crown domains line an extended DNA channel, with the wing domain spanning outwards (**Fig. 3 B**). The diameter of the central channel varies from ~35 Å at the crown domain, to ~33 Å in the clip domain, and ~22 Å in the tightest part of the stem (**Fig. 3 A**). This is sufficiently wide for translocating B-form DNA (58). The local resolution of the portal protein **(Fig 3 C)** shows the wing domain particularly well resolved, with lower resolution for the crown and especially the clip domain. A reconstruction of the portal protein from a separate sample of HK97 prohead II. featuring the *in situ* portal protein, but without terminase (unpublished data), presents an informative comparison **(Fig. 3 D**). While the global resolution of this ‘empty prohead’ portal is lower, at 4.1 Å, low-pass filtering each map to the same threshold indicates a distinct improvement in the definition of the clip domain, where density corresponding to α-helices can be clearly distinguished. Further differences appear also in the crown domain, which is extended in terminase absence.

**Figure 3:**
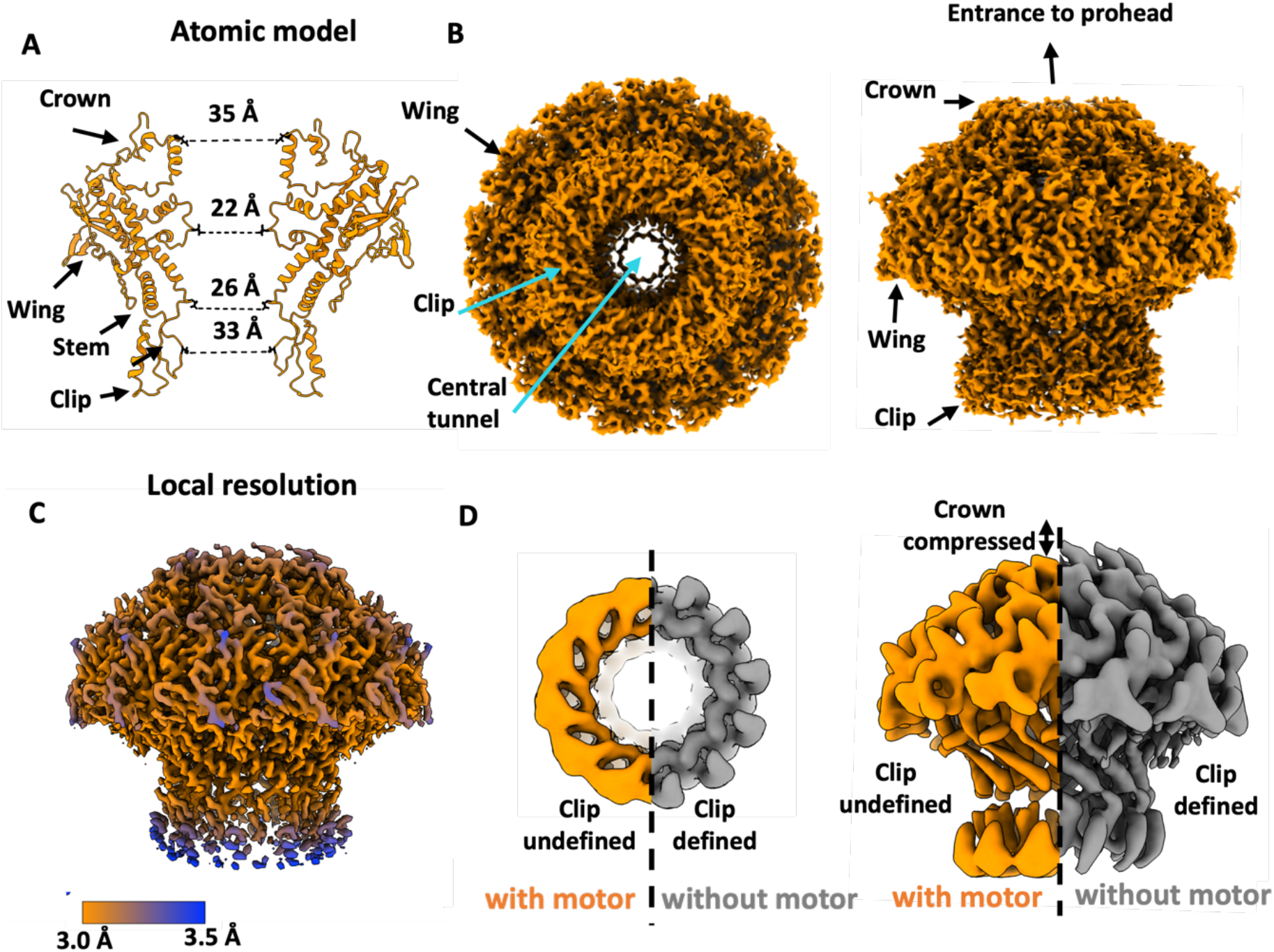
Structure of the portal protein. **A** to **C** - C reconstruction of the in-situ portal protein at 2.98 Å resolution. **A** – ribbon diagram with only two opposing subunits. **B,C** – portal density. **D** – comparison of portal protein maps in the presence and absence of the motor, viewed from the clip (left) and along the tunnel axis (right).

### Reconstruction of the packaging motor comprising the portal-large terminase complex

In spite of attempts to fix the large terminase motor in a single state—both by introducing a *cos* site in the DNA substrate and providing a high concentration of ATP*γ*S— processing of the data indicates that the motor is highly flexible and heterogeneous. Icosahedral symmetry expansion of the capsid followed by focussed 3D classification produced two broad portal/large terminase classes. Both showed strong signal at the large terminase locus but little definition, indicative of flexibility (**Fig. S 2 B, C**). Indeed, further 3D classification revealed a whole spectrum of conformation, only several of which displayed defined subunits (**Fig. S 2 C**). Subsequently, the terminase oligomer reconstruction represents just 17,584 particles out of a total of approximately 82,279 portal/large terminase particles.

The asymmetric reconstruction, derived at 8.8 Å resolution (**Fig. 4 A**), allows the five large terminase subunits to be clearly observed, which are arranged in a pentameric ring encircling the dsDNA substrate. This stoichiometry for large terminase is in keeping with previous fluorescent photobleaching experiments (43) and the motor assemblies of other dsDNA bacteriophages (2,4–6,59). When the same particles were re-extracted to encompass both the large terminase and the portal protein in complex, the resolution improved to 7.4 Å (**Fig. 4 B**), consistent with a more structurally rigid conformation. Clearly resolved rod-like density for the DNA within the lumen of the large terminase appears to make contact with both the N- and C-terminus domains of large terminase monomers in both reconstructions (**Fig. 4)**.

**Figure 4:**
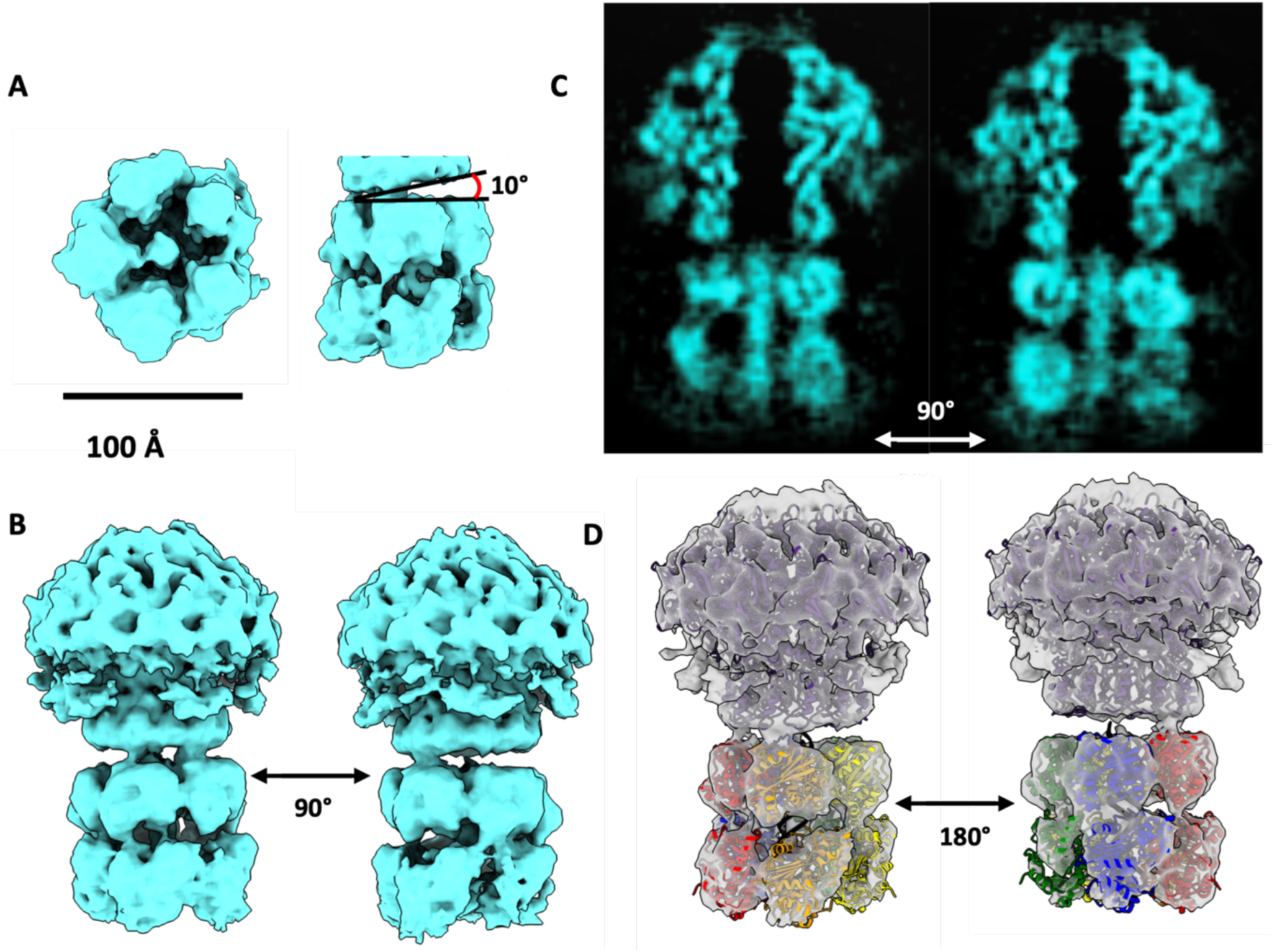
Asymmetric reconstruction of the HK97 DNA packaging motor. **A** - pentameric motor surrounding DNA substrate. **B** - the portal-terminase motor complex. **C** - cross sectional views of the portal-motor complex. **D** - result of the rigid body docking of terminase domains, derived from the crystal structure of the HK97 large terminase, into the EM density with the fitted models shown as ribbon diagrams coloured by subunit.

The large terminase reconstruction is in good agreement with the crystal structure of the monomer (43) with densities for each ATPase and nuclease domain well resolved (**Fig. 4 A,B,C**). Flexibility within the complex is likely confined to inter-domain and inter-subunit movement, as opposed to domain rearrangement, consistent with the globular nature of the two domains and the ability to crystallise single-domain constructs of related large terminases more readily than full-length constructs (60–66). Thus, individual domains were subject to rigid-body fitting into the density (55) (**Fig 4 D**). The more compact nuclease domains were fitted into the ring directly below the portal, with the N-terminal ATPase domains positioned more flexibly beneath, with 25 base pairs of DNA spanning the central tunnel of the terminase assembly.

## DISCUSSION

### Motor assembly induces asymmetry in the portal protein

The density corresponding to the portal clip domain within the packaging complex, compared to the portal clip domain in the empty prohead, is poorly resolved. This suggests distortion away from C_12_ symmetry during packaging. We hypothesise that the plasticity of the clip domain compensates for the symmetry mismatch at the portal-large terminase interface and facilitates interactions with the large terminase. Meanwhile, the reduced height of the crown domain in the portal/large terminase complex echoes structural changes in related portal proteins, where the mounting pressure from DNA within the capsid is thought to induce portal compression (34,37). Flexibility within each of these domains has been indicated in studies of other viruses during DNA packaging (37,67).

### Small terminase likely dissociates after inducing large terminase endonuclease activity

Small terminase is essential for the stalling of the HK97 large terminase motor, and thus present in the sample, but no obvious density corresponding to a small terminase is apparent within the density maps. The nonameric ring of small terminase has a molecular weight of 160 kDa, which should be discernible even if positioned flexibly. This indicates a transient role for small terminase in the complex formation. We hypothesise that the small terminase remains bound to the *cos* site throughout packaging, acting as a roadblock for incoming large terminase, and instigating the conformational change required to switch the motor from displaying ATPase activity to endonuclease activity. In turn, small terminase may dissociate from the DNA (**Fig. 5C**). The documented stimulatory effect of the small terminase (43), in this model, would then be attributed solely to recruitment of large terminase to the viral DNA, thus enhancing the number of packaging initiation events.

**Figure 5.**
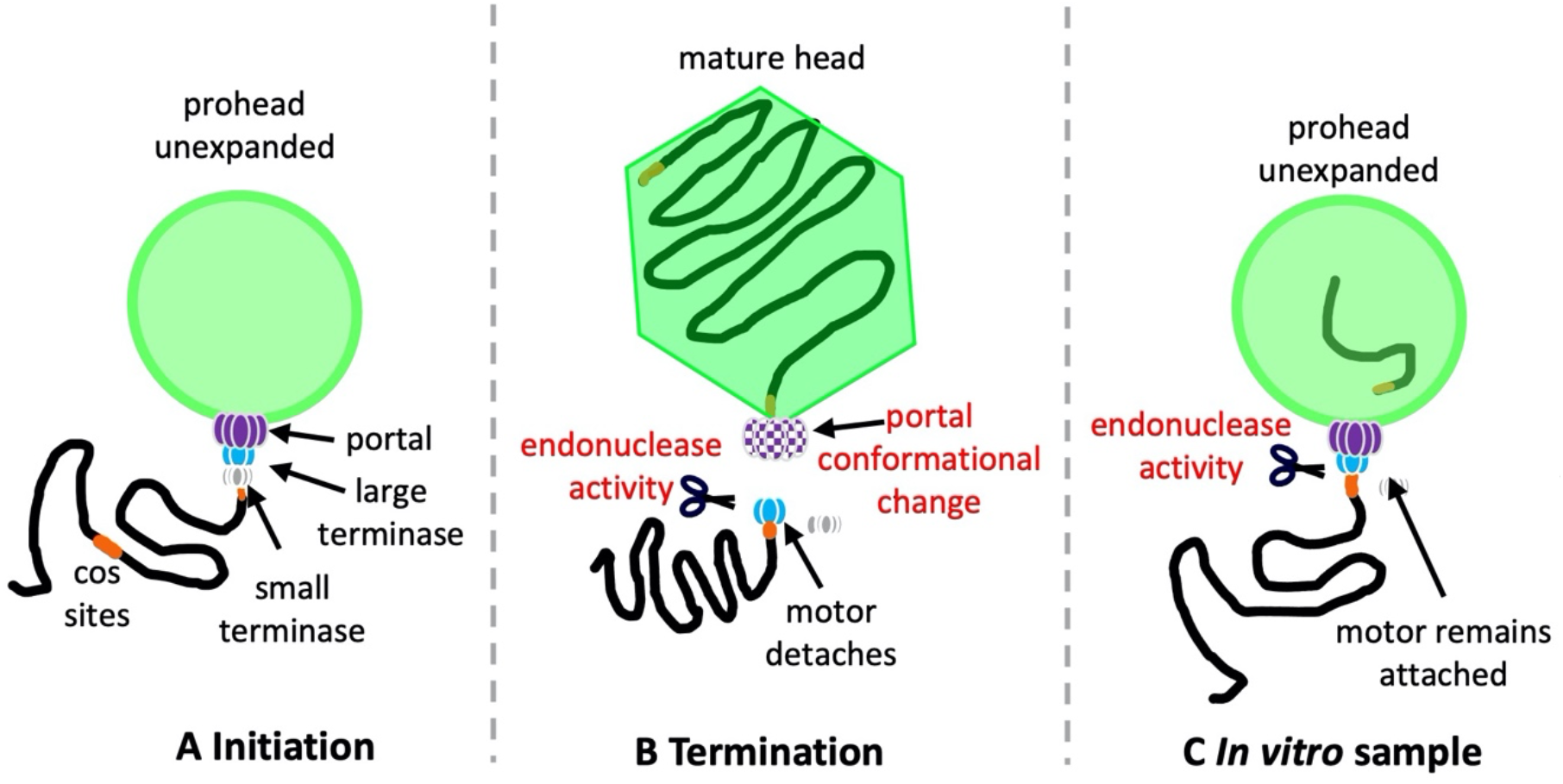
Packaging termination in vivo and in vitro. **A** - Initiation complex of HK97 DNA packaging in vivo. **B** - Termination complex of HK97 packaging in vivo. **C** - HK97 stalled packaging complex present in vitro, after cleavage of DNA with the large terminase assembly remaining attached.

### DNA cleavage at the large terminase - portal interface mimics termination

One striking feature of the portal-large terminase map is the apparent lack of DNA density within the portal channel (**Fig. 4 C**). DNA extends through the lumen formed by the large terminase oligomer and stops at the interface between the large terminase and portal. The length of this DNA is consistent with approximately 25 bp of B-form DNA. Packaging assays consistently show protected DNA within proheads of ~1.3 kbp, suggesting that the motor has packaged from the 5’ end of the DNA substrate and paused at the *cos* site (**Fig. 1 A**). This mimics the packaging termination state: on reaching a *cos* site after packaging one complete genome unit, large terminase endonuclease activity is activated, the DNA is cleaved, and the motor dissociates (**Fig. 5 B**). In our sample however, in spite of clear DNA cleavage, large terminase remains bound to the portal. This is also consistent with the docked crystal structure of the large terminase, which places the nuclease domains next to the cleaved DNA end near the portal tunnel entrance (**Fig. 4 D**).

This suggests that packaging termination may occur *via* two steps, controlled by different signals: (1) DNA cleavage at a *cos* site is mediated by small terminase large terminase interactions; (2) release of the large terminase occurs in response to headful pressure signalling, likely relayed through the portal. In the structure of prohead II, presented here, the capsid remains relatively empty, and unexpanded, as the length of DNA inside of ~1.3 kb represents just ~3% of 39.7 kb genome. In the absence of the headful signalling the large terminase hence does not detach, but cleavage at the *cos* site can still occur (**Fig. 5**).

### Translocation is mediated by interdomain extension-contraction

An asymmetric interaction between the large terminase and the portal clip domain is apparent in the portal-large terminase reconstruction (**Fig. 4**). Clear contact with the portal protein appears only between one large terminase subunit, with a second adjacent subunit showing limited interactions. The entire large terminase assembly is also tilted 10 ° to the portal tunnel axis (**Fig. 4A**). Asymmetric alignment was similarly observed for the Φ29 motor, where the large terminase channel is tilted at 12.5° to the portal axis (59). However, in contrast, the Φ29 portal-terminase interaction is mediated by a pRNA, which is absent in the HK97 system, while the terminase itself lacks nuclease activity (68).

In the docked crystal structure of the HK97 large terminase (**Fig. 4 D**), the two subunits which contact the portal (depicted in red and orange), are separated axially by a further ~4.5 Å (or 1.5 base pairs of DNA), relative to the subunits shown in green and blue (69). Both domains of the extended subunits contact DNA (**Fig 6 B)**, whilst the contracted subunits contact DNA via the nuclease domain only (**Fig 6 B).** Experimental data on multiple related large terminase has shown that ATP bound subunits grip DNA where ADP bound subunits do not (7,70–72); and so, by analogy the extended HK97 subunits may be assigned ATP*γ*S bound whilst contracted subunits can be assigned ADP bound. The presence of ADP in the sample would require slow hydrolysis of ATP*γ*S, as observed in DNA translocases (73). Notably, the contracted subunits display broken contact with the clip domain of the portal protein, which could potentially facilitate a cyclically changing pattern of terminase-portal interactions throughout packaging. The final large terminase subunit, shown in yellow, shows a less dramatic extension of 2 Å, which could correspond to an ATP*γ*S hydrolysis transition state (**Fig 6 A**) (73). An overlay of each docked large terminase ATPase domain after aligning docked nuclease domains, depicts the interdomain shifts around the pentameric ring (**Fig 6 C**). As such, contraction and relaxation of subunits is likely involved in DNA translocation – with ATP hydrolysis coupled to subunit contraction which pushes the DNA substrate into the prohead.

**Figure 6:**
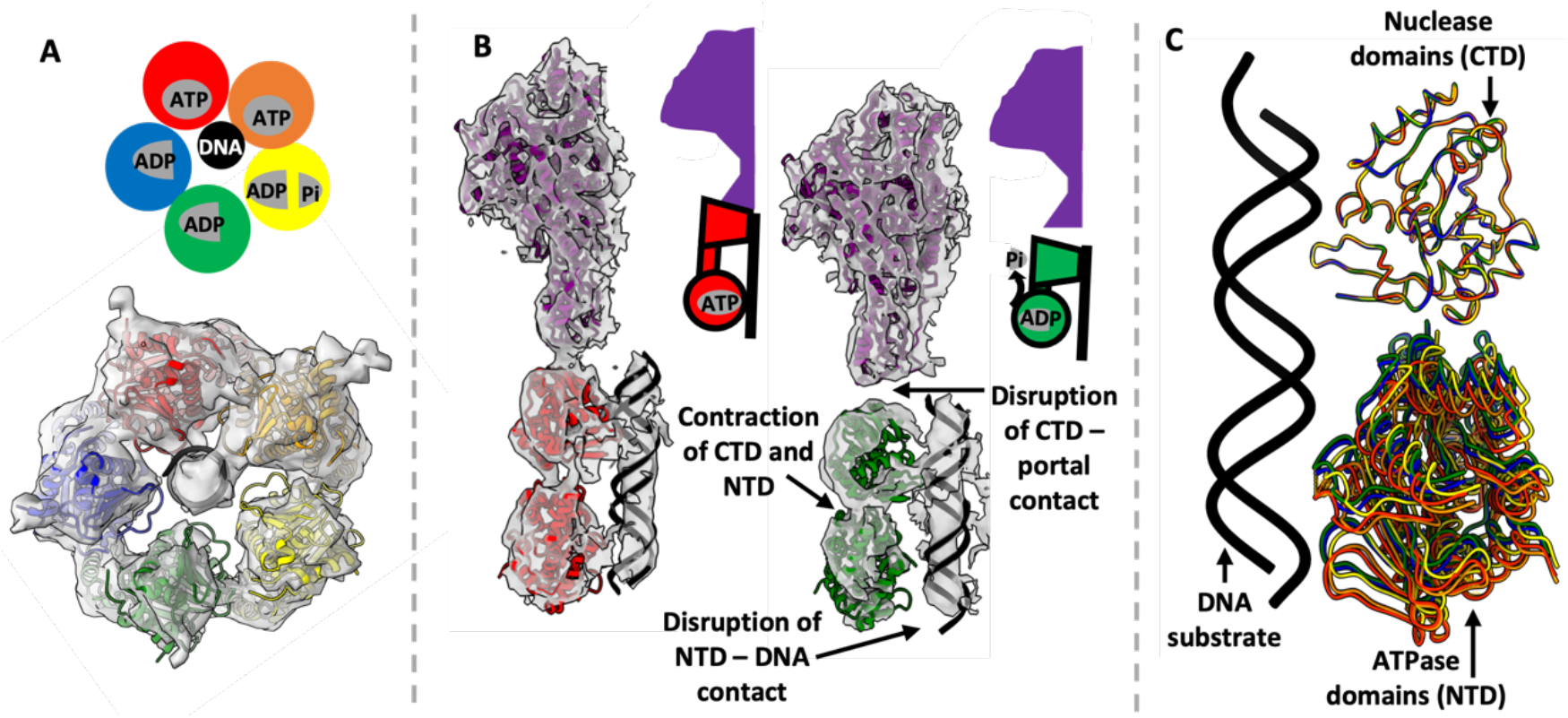
Structural comparison of HK97 large terminase monomers of the motor. **A** - proposed nucleoside binding pattern. **B** - comparison of proposed ATP and ADP bound subunits. **C** - overlay of large terminase monomers after aligning the nuclease domain

Contraction of large terminase subunits from an extended ATP bound state, to a contracted ADP bound state, have also been proposed as translocation mechanisms of the bacteriophages T4 and Φ29 (4,59). For T4, ATP hydrolysis is proposed to occur sequentially around the ring, so that only a single subunit is ever present in the ‘tense’ contracted state (74). This does not entirely agree with the HK97 structure where subunits display a range of conformations. Meanwhile, the five bacteriophage Φ29 ATPase domains are thought to make sequential shifts toward the vestigial nuclease domains (which remain in a planar ring throughout). This completes a cracked helix to planar transition, in four ATP hydrolysis steps (59). Transposition of this mechanism onto our proposed HK97 model is also problematic since transition through full ADP occupancy could destabilise the terminase-portal contact.

An alternative comparison can be made with the DNA translocation mechanism proposed for the *E. coli* translocase FtsK (73). In this mechanism, a molecular machine comprises six subunits, with each adopting a unique conformation with variable angles of separation between the α and β domains (as opposed to extension) (**Fig. 7C**). Three subunits bound to ATP*γ*S, and a fourth trapped in ATP*γ*S hydrolysis, display DNA binding residues in a spiral. The remaining two subunits are ADP-bound and do not engage with DNA (73). The proposed translocation mechanism for FtsK never passes through a fully ADP bound state, with the rolling change in conformations around the ring shifting sequentially, akin to a spiral escalator (73). This mechanism is more consistent with the HK97 structure: where subunits display a range of conformations and DNA is also engaged with the subunits in extended conformation, assigned as ATP*γ*S bound. A comparison between the individual monomers of HK97, Φ29 and FtsK ATPases **(Fig. 7)** shows how each system depends on a varied extension of individual subunits mediated by interdomain movements.

**Figure 7:**
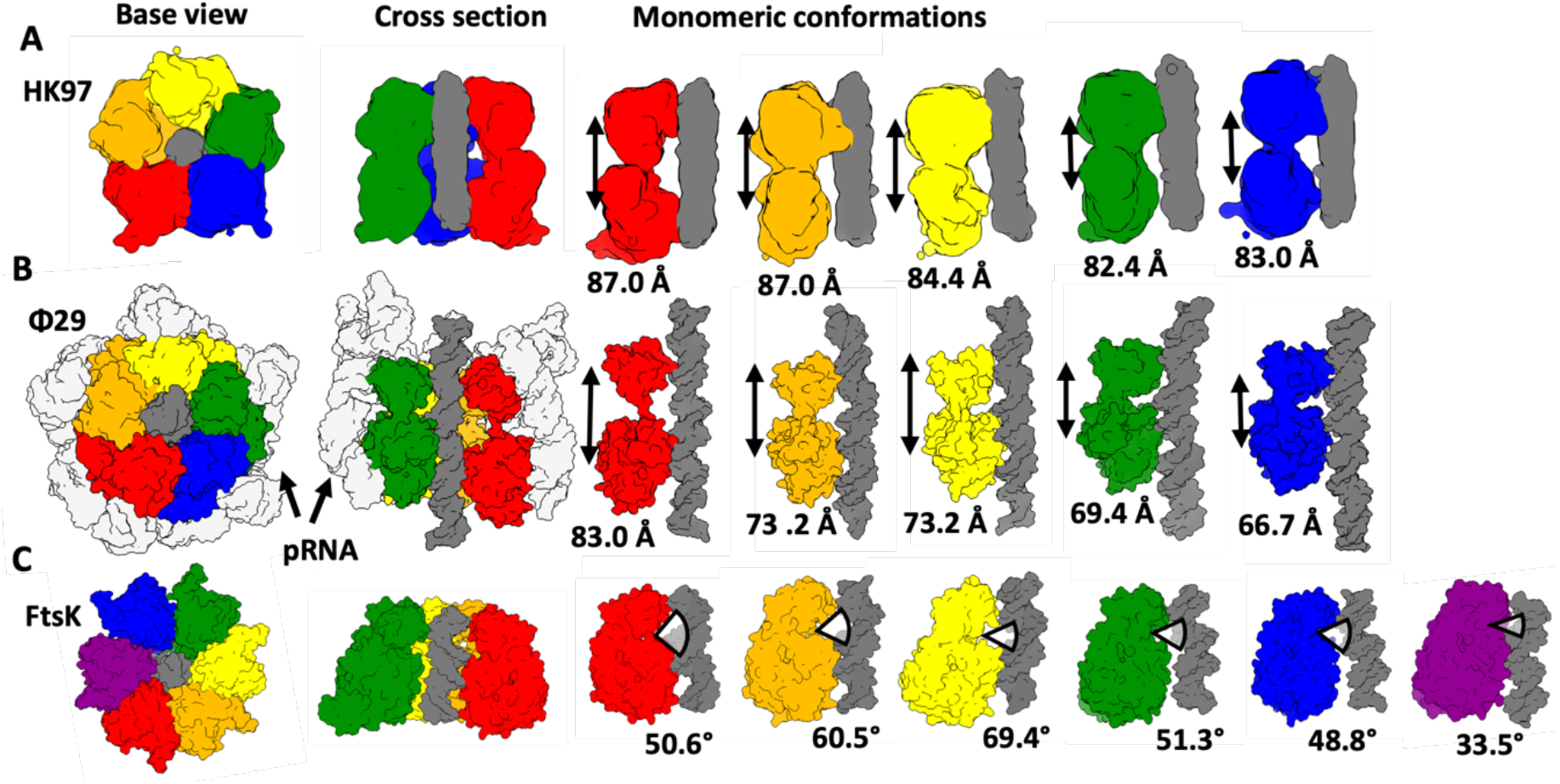
Comparison of domain adjustments in HK97, Φ29 and Ftsk DNA translocation motors. Subunits are coloured differently and shown as molecular surfaces. DNA (Φ29 and Ftsk) or DNA density (HK97) is in dark grey. For each motor (**A** – HK97, **B** - Φ29 and **C** – Ftsk) two orthogonal views of the assembly are on the left, and individual subunits along with DNA are shown on the right

### Conclusions

Here we present a structure of the complete packaging machinery of HK97 determined by cryo-EM. The high-resolution structure of the prohead shows that the capsid adopts the immature, unexpanded state (prohead II), present at the initiation of DNA packaging. However, the well-defined DNA density within the large terminase pentamer ends abruptly at the portal protein interface, characteristic of DNA cleavage seen at packaging termination. This suggests that packaging termination requires two distinct signals, and while the *cos* sequence present in the DNA substrate is sufficient to induce DNA cleavage by the large terminase, headful pressure is required for motor dissociation. The cryo-EM structure of the portal protein, derived at 2.98 Å resolution, shows that the clip domain deviates from the C_12_ symmetry, facilitating interactions with the five subunits of the large terminase. Docking of the large terminase subunits into the motor map suggests that the observed extension and contraction between the two domains of each subunit is involved in ATP-driven translocation of DNA into prohead, with both the N- and C-terminal subunits making contacts with DNA (55). This translocation model has commonalities with mechanisms proposed for the Φ29 and T4 packaging motors (59,74), as well as the DNA translocase Ftsk (73), each of which is proposed to utilise inter-subunit contraction to move DNA through the central pore of an oligomeric ring. In particular, the rolling ATP-ADP exchange of FtsK and variable domain arrangement fits well with the highly asymmetric nature of the HK97 large terminase pentamer, and the limited interaction with the portal protein.

## DATA AVAILABILITY

Atomic coordinates for the prohead and portal protein and maps have been deposited with the Protein Data Bank and Electron Microscopy Data Bank under accession numbers 8CFA/EMD-16624 and 8CEZ/EMD-16614 respectively. Maps for the isolated large terminase complex (**Fig 4 A**) the portal/large terminase complex (**Fig 4 B,C,D**), and the whole packaging complex (**Fig 1 D**) have been deposited to the Electron Microscopy Data Bank under accession numbers EMD-16654, EMD-16653 and EMD-16649 respectively.

## ACKNOWLEDGEMENT

The authors would like to thank Turkenburg J., Hart S. and Blaza J.N. for assistance in cryo-electron microscopy at York and to Morris K.L. for data collection assistance at eBIC. We would also like to acknowledge Chechik M. for assistance in protein production and to thank Hesketh E.L., Thompson R.F. and Maskell D.P. for facilitating the collection of additional data sets and help with processing. Bacterial strains and HK97 mutant strains used in procapsid production were provided by Duda, R. L. Lastly, thanks to Duda, R. L. and Jardine,P.J. for helpful discussion and assistance setting up the HK97 packaging system.

---

Table 1. The quick brown fox jumps over the lazy dog.

Figure 1. The quick brown fox jumps over the lazy dog.

Figure 1: Structure of a complete HK97 packaging complex.

Figure 3: Structure of the portal protein

Figure 2: Icosahedral reconstruction of the HK97 prohead II at 3.06 Å resolution

Figure 4: Asymmetric reconstruction of the HK97 DNA packaging motor

Figure 6: Structural comparison of HK97 large terminase monomers of the motor

Figure 7: Comparison of domain adjustments in HK97, Φ29 and Ftsk DNA translocation motors

Figure S2: Data processing schemes

Table S1: Cryo-EM data collection parameters for stalled HK97 packaging complexes

Table S3: Model refinement statistics for the prohead and portal protein

## SUPPLIMENTARY INFORMATION

**Table S1:** Cryo-EM data collection parameters for stalled HK97 packaging complexes.

|  |  |
| --- | --- |
| Location | eBIC |
| Grid type | R 3.5/1 with Ultrathin Carbon |
| Microscope | Krios 2 |
| Detector (Mode) | K3 (counting super-resolution) |
| Accelerating Voltage (kV) | 300 |
| Spherical aberration / mm | 2.7 |
| Calibrated pixel size (Å) | 1.34 |
| Nominal Mag. | 64,000x |
| Total dose (eÅ<sup>-2</sup>) | 40 |
| Nominal defocus range (μm) | -0.5 to -2.5 |

**Figure S2:**
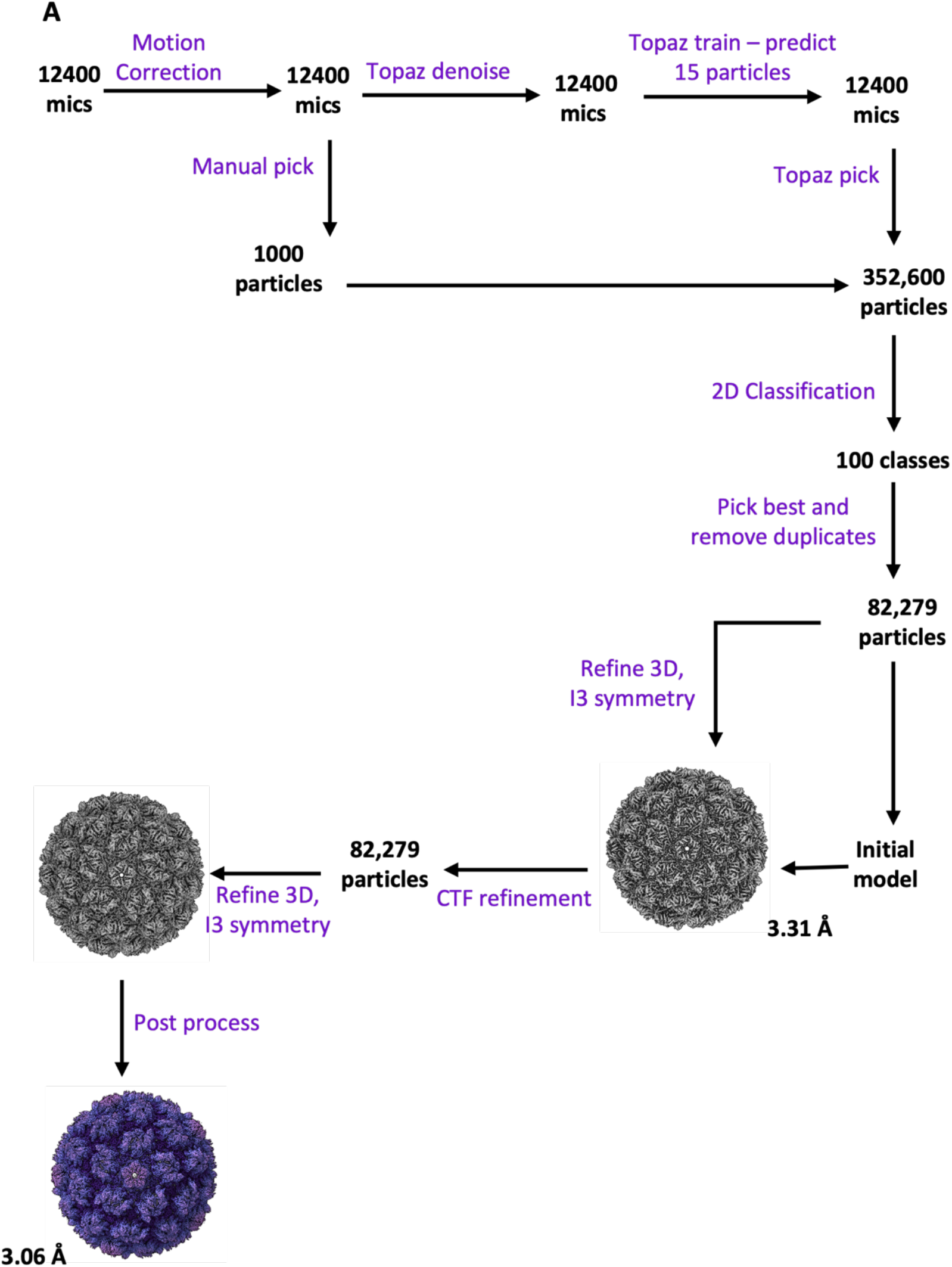
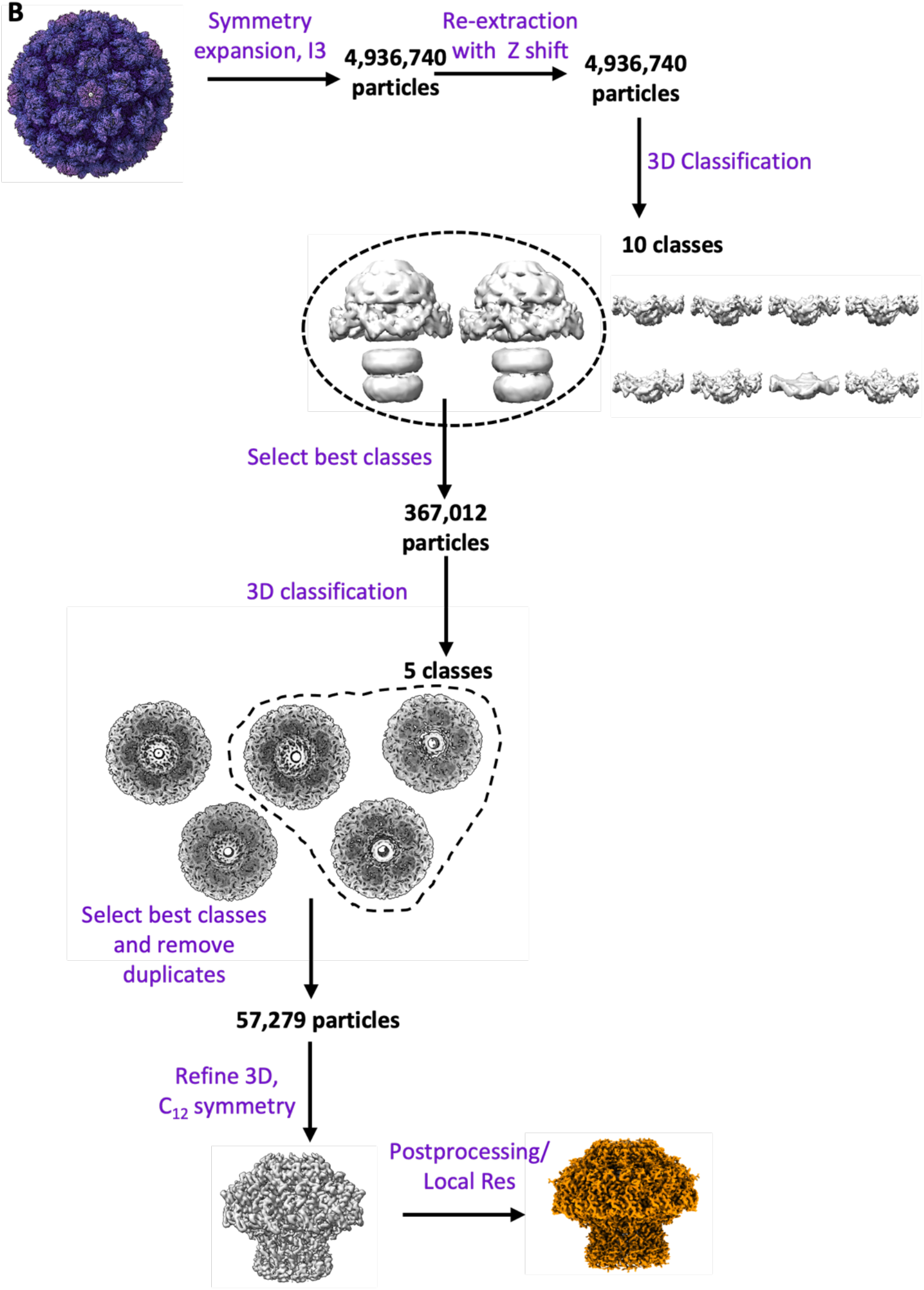
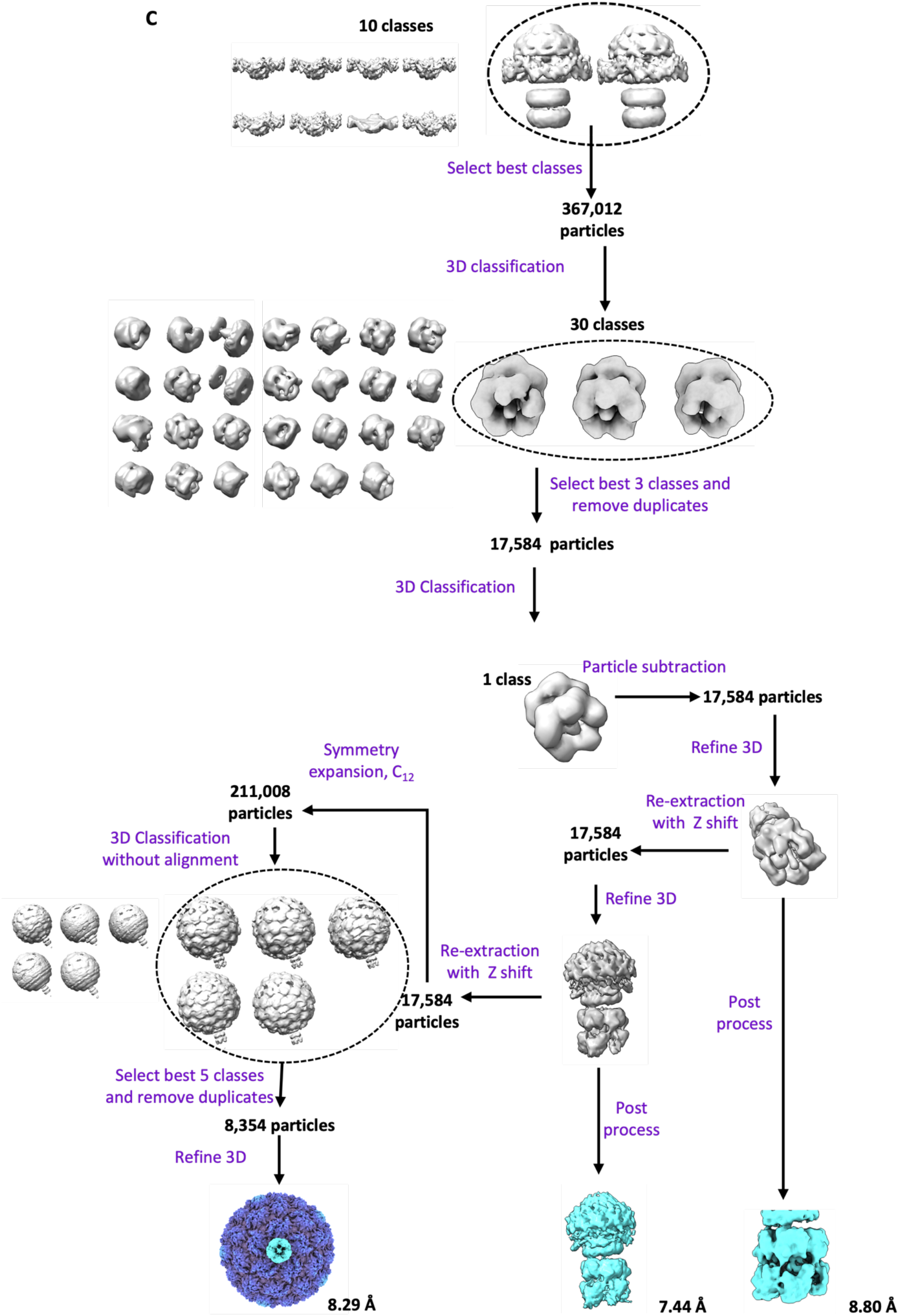
Data processing schemes for **A** - the HK97 prohead, **B** - the HK97 portal protein and **C** - the HK97 large terminase assembly, portal/terminase complex and complete packaging complex comprising prohead II, portal protein and large terminase.

**Table S3:** Model refinement statistics for the prohead and portal protein.

| **Prohead** |  |
| --- | --- |
| Symmetry | I3 |
| Resolution (FSC 0.143) | 3.06 |
| Map-to-model correlation | 0.80 |
| MolProbity score | 1.53 |
| RMS deviations |  |
| Bond lengths / Å | 0.005 |
| Bond angles / ° | 0.9 |
| Ramachandran plot / % |  |
| Favored | 95.4 |
| Allowed | 4.5 |
| Outlier | 0.1 |
| **Portal protein** |  |
| Symmetry | C12 |
| Resolution (FSC 0.143) | 2.98 |
| Map-to-model correlation | 0.68 |
| MolProbity score | 1.9 |
| RMS deviations |  |
| Bond lengths / Å | 0.003 |
| Bond angles / ° | 0.6 |
| Ramachandran plot / % |  |
| Favoured | 91.6 |
| Allowed | 8.1 |
| Outlier | 0.3 |

